# Active and Probe-Free Intracellular Rheology via Phase-Sensitive Thermoviscous Flows

**DOI:** 10.1101/2025.04.07.647540

**Authors:** Iliya D. Stoev, Madison Bolger-Munro, Antonio Minopoli, Susan Wagner, Venkat R. Krishnaswamy, Elena Erben, Kai Weißenbruch, Nicola Maghelli, Martin Bastmeyer, Carl-Philipp Heisenberg, Moritz Kreysing

## Abstract

Determination of the rheological properties of cells is known to require active measurements, which largely depend on the internalisation of mechanical probes. Here we circumvent this problem via the introduction of Rheo-FLUCS: an active, yet probe-free approach that leverages light-induced flows to access the mechanics of complex systems. While Rheo-FLUCS is facilitated by thermoviscous expansion phenomena rather than external forces, we here show equivalence in its ability to measure canonical viscoelastic properties. Specifically, we demonstrate a phase-lag equivalence with probe-dependent active microrheology in a wide range of physically different, yet chemically identical materials. We exemplify the utility of Rheo-FLUCS in three distinctly different biological systems: compound-treated mouse fibroblasts (NIH-3T3), genetically modified human osteoblasts (U2OS) to elucidate the role of myosins in cytoplasmic mechanics, and early ascidian oocytes of *Phallusia mammillata* at fertilisation stage. This probe-free methodology represents a transformative advance in cellular rheology, enabling non-invasive and precise mechanical measurements across diverse biological systems without perturbing their natural state through probe internalisation.

## Main

Important roles of mechanics become increasingly recognised in the life sciences, and specifically in cell and developmental biology across scales, ranging from force-generating molecules,^1,2^ through condensates with functional material properties,^3–5^ to ooplasm and embryonic mechanics.^6–8^ Passive rheological measurements may be probe-free, but are known to be insufficient in measuring the mechanics of many out-of-equilibrium systems and active matter, such as living systems. The limited thermal motion of a probe poses an operational limit to the attainable viscoelastic range, and the generalised Stokes-Einstein relation, which bridges particle dynamics with material properties, breaks down.^9^ Active measurements, on the other hand, are capable of yielding faithful results even within living cells, but usually require the internalisation of a mechanical probe,^10,11^ which might be difficult to achieve or report non-physiological responses. Active measurements that are probe-free would therefore combine the advantages of two disparate approaches.

Previously, it has been suggested that thermoviscous flows,^12,13^ the physical mechanism underlying Focussed Light-Induced Cytoplasmic Streaming (FLUCS)^14–19^ can be used to assess the mechanical properties of the cytoplasm by quantifying the flow displacement upon application of a driving stimulus.^20^ However, it remained to be shown that thermoviscous flows can also be used to extract canonical rheological parameters, such as the phase angle, as a quantitative metric of viscoelastic material behaviour in complex, non-Newtonian fluids on the microscale.^21–26^ Leveraging the optical control capabilities of FLUCS, we advance the flow perturbations, yielding a new phase-sensitive microrheology method called Rheo-FLUCS, which provides access to intracellular rheology in an active, yet probe-free way, thereby significantly advancing the fields of materials science and mechanobiology.

## Results

### Rheo-FLUCS: phase-sensitive active microrheology enabled by thermoviscous flows

Previously, it has been demonstrated how the rapid scanning of an infrared laser beam resulted in localised and directed flows within fluids of arbitrary viscosity, including the cytoplasm of a worm embryo or that of yeast cells.^13,14,19,20^ These thermoviscous flows arise due to the combined effect of temperature-induced viscosity changes and thermal expansion of the fluids.^12,13^ Periodically reversing the direction of laser scanning generates oscillatory flows, the amplitude of which was used to determine relative mobility of the cytoplasm as a proxy for its rheology.^14^

Here, we conceptually extended the measurements to collect temporally resolved data, specifically from measurements that are highly synchronised with the laser-induced stimulus. This allows quantitative signal processing and access to phase-sensitive complex amplitudes of the oscillation, after appropriate Fourier-domain filtering.

In classic bulk rheology,^27–29^ stress-strain relations are widely used to characterise viscoelastic materials. For this, a periodic external force is used to induce oscillating stresses with angular frequency *ω*, which in turn produce an oscillatory shear deformation, with the same frequency but a temporal delay called ‘phase lag’. Stress that is strictly in-phase with the strain (phase angle *φ* ∼ 0°) signifies a purely elastic Hookean solid with finite frequency-dependent storage modulus *G*^′^ (*ω*) and comparably negligible frequency-dependent loss modulus *G*″ (*ω*). On the other hand, a fully out-of-phase stress-strain relation (*φ* ∼ 90°) corresponds to a purely viscous Newtonian liquid with finite frequency-dependent loss modulus *G*″ (*ω*) and negligible storage modulus *G*^′^ (*ω*). In-between these two extremes, the entire complex viscoelastic regime for non-Newtonian fluids is marked by phase angles 0° < *φ* < 90°, and specifically the relation 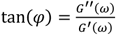 is frequently used to practically access the phase angle, as a relative measure of viscous and elastic properties of a material.

In this work, we suggest that rather than requiring the use of an internalised force probe^30–32^ to induce stresses that strain a material, flows that deform a medium can directly be induced via thermoviscous laser actuation, while still maintaining the quantitative phase relation between the driving stimulus and the material response (Figure 1a). To this end, we directionally and repeatedly translate an infrared laser spot across the sample with a frequency in the low-kilohertz range, thereby inducing thermoviscous flows within it. The straining caused by this localised stimulus is monitored through the motion of a discernible feature within the sample, *e*.*g*. a tracer particle, which upon periodic switching of the scan direction reports on the sample response in the form of an observable oscillation (Figure 1b).

**Fig. 1.**
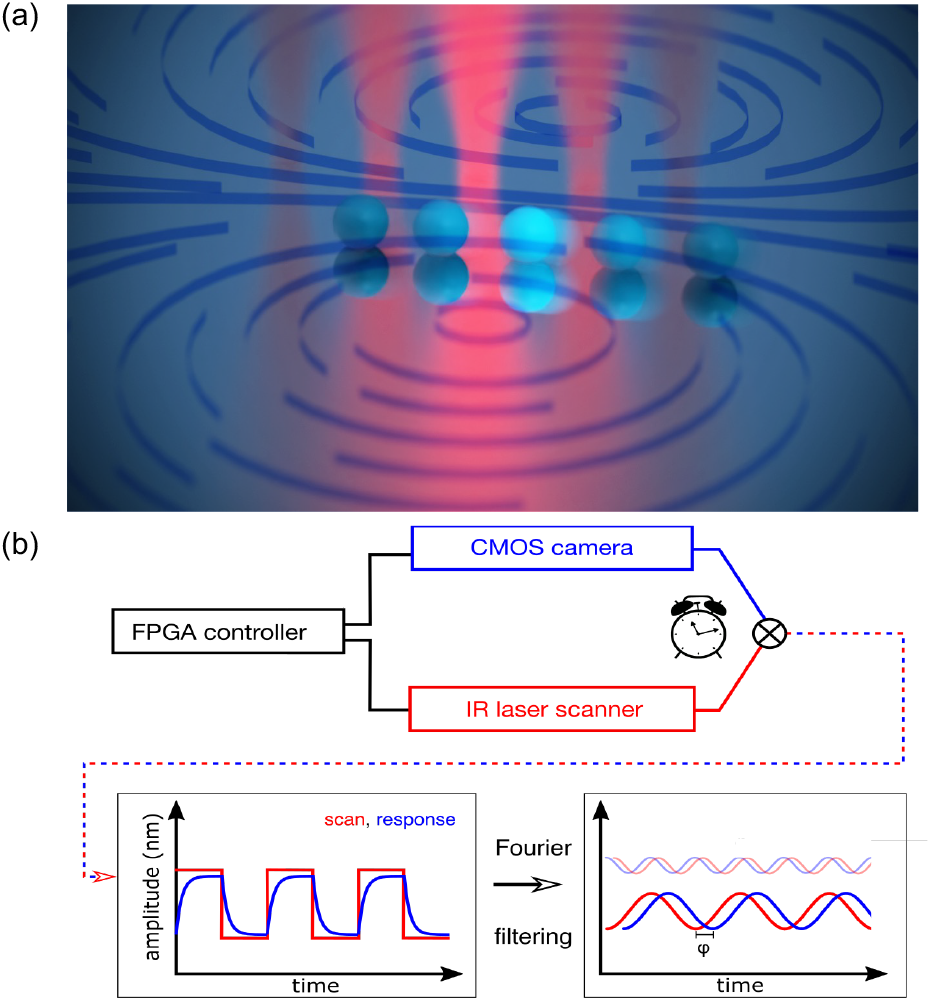
Conceptual and technical drawings of the phase-sensitive microrheology pipeline used by Rheo-FLUCS to extract rheological properties. (a) Conceptual representation of the laser scanning (flow stimulus in red) as active oscillatory driving and the resulting time-delayed light-induced flows reported by a tracer (sample response in cyan). (b) Technical implementation of the microrheology pipeline that relies on synchronous triggering of the camera and the laser by a central controller. Utilising this fine synchronisation, we attributed any phase lag between the driving stimulus and the sample response to the material properties of the sample. This allows dynamic tracking of mechanical information in real time, where Fourier filtering eliminates passive and active noise present at all frequencies other than the excitation.

We argue and show that despite using an alternative stressor to conventional rheology, the phase angle, as canonical quantifier of fluidity, remains the same. We validate this hypothesis across a wide range of physically different, yet chemically identical systems established by Charrier *et al*.^33^ (Figure 2a). These allow the independent tuning of elastic (*G*^′^ (*ω*)) and viscous (*G*″ (*ω*)) contributions in well-characterised environments.

**Fig. 2.**
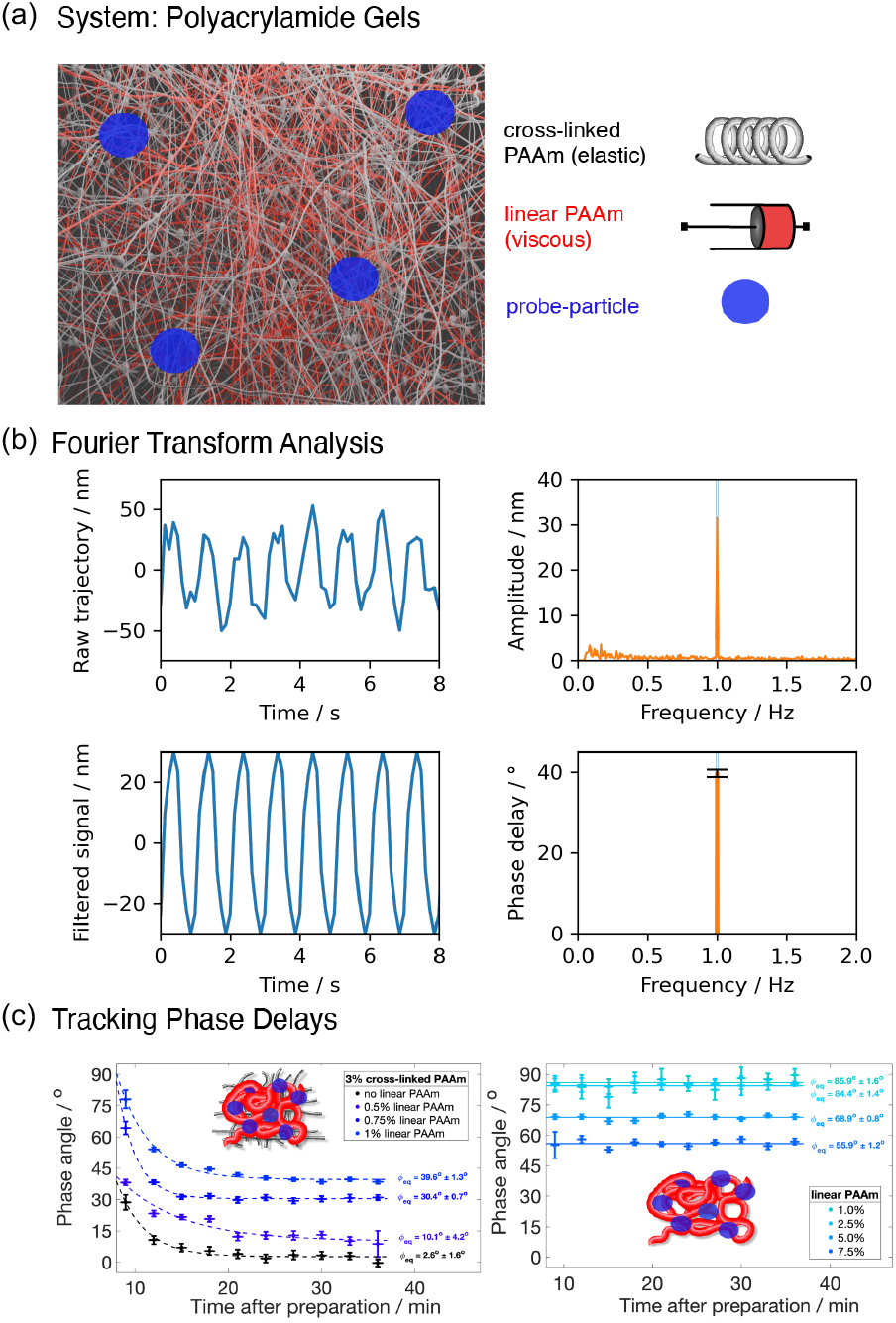
Viscoelastic model system, data extraction pipeline and sensitivity to the time-resolved solidification of fluids. (a) Tunable polyacrylamide gels as model viscoelastic systems: independent tuning of the elastic and viscous contributions is achieved through varying the concentrations of cross-linked (grey) and linear (red) polyacrylamide, respectively. Oscillations are flow-induced, independently of mechanical probes. Small tracer particles merely report on the oscillations and are not necessarily required in intrinsically heterogeneous living systems. (b) Fourier filtering analysis is used to convert a noisy raw trajectory into low-noise canonical response of the material, where the clean signal is reconstructed by filtering for a given oscillation frequency. This reconstructed signal provides access to the phase delay between the driving stimulus and the sample response, which reveals the mechanical properties of the test sample (data shown, 3%vol cross-linked PAAm and 1%vol linear PAAm probed at 1 Hz). (c) Real-time tracking of gelation in cross-linked polyacrylamide gels (left) and entanglement in liquid suspensions of linear polyacrylamide chains (right). The dashed curves represent exponential fits with matching relaxation times for gelation (mixtures of cross-linked and linear PAAm), while solid curves confirm the relatively constant repeated measurements in viscoelastic fluids.

### Real-time tracking of mechanical properties in model viscoelastic systems

First, we asked if Rheo-FLUCS can quantitatively measure the mechanical behaviour of Newtonian liquids and report their chemically induced solidification. For this, we started by preparing dilute (highly fluid) linear polyacrylamide (PAAm) suspensions and oscillated 1-*μ*m fluorescent polystyrene tracer beads to visualise the straining. Using a custom developed Fourier-transform decomposition tool (Figure 2b), we repeatedly extracted the same phase angle (Figure 2c), with a value of *φ* around 85.91° ± 1.59°, close to a perfect liquid with virtually no elastic component.

Next, we investigated if the method could reliably report the real-time formation of a cross-linked gel (Figure 2c, black data, 3%vol PAAm, with monomer-to-bis-acrylamide cross-linker ratio of 100:1). The initial value of *φ* close to 30° signified the onset of gelation and we observed a progressive drop in the phase angle until a steady state was reached 20 min after sample preparation at around 2.63° ± 1.55°. These results so far show that Rheo-FLUCS is able to qualitatively report the dynamic solidification of a fluid, with a terminal phase angle characteristic of Hookean, perfectly elastic solids.

### Quantitative validation of phase angles using a force-driven approach

To confirm the equivalence between phase angles extracted from flow- and force-driven microrheology experiments, we compared a range of physically distinct, yet chemically near identical materials in their responses to both types of rheology. Specifically, in the force-driven microrheology we measured oscillatory step-stress perturbations via a magnetic force probe.

To start, these measurements confirm that for both extremes, *viz*. the predominantly fluid PAAm monomers and solidified PAAm gels, Rheo-FLUCS yields also quantitatively the same responses as force-driven microrheology (Figure 3).

**Fig. 3.**
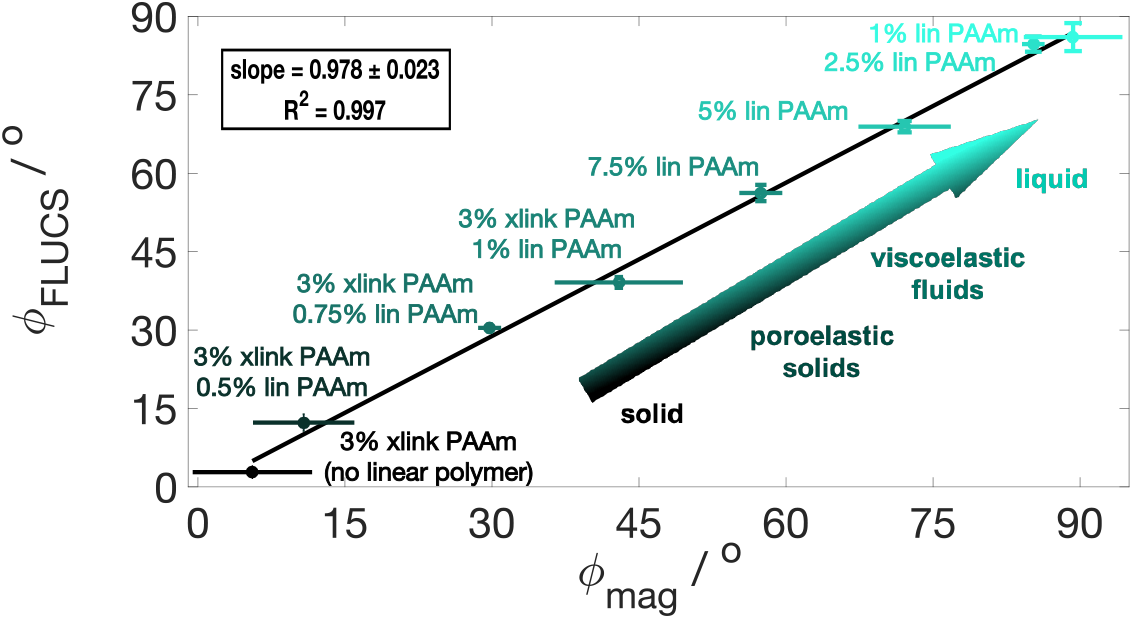
Equivalence of phase angles extracted via Rheo-FLUCS and force-driven rheology measurements. Direct comparison between phase lags obtained from Rheo-FLUCS and magnetic force driving for chemically identical, yet physically different mixtures of cross-linked (xlink) and linear (lin) polyacrylamide (PAAm). The data points represent weighted averages of five independent measurements. The linear fit confirmed the hypothesis that the stress-strain formalism relating phases and viscoelasticity from conventional bulk rheology offers a valid analytical treatment of data generated optofluidically with Rheo-FLUCS.

Next, we assessed if also in the intermediate regime of viscoelastic materials, a direct linear proportionality for flow- and force-based measurements holds true. For this, we polymerised PAAm monomers into linear chains. In the absence of cross-linkers, this yields highly viscous solutions, with elasticity increasing with monomer concentration (from 2.5%vol to 7.5%vol) and phase angles decreasing down to about 60°, in agreement with the force-based reference measurements. This material response can be attributed to polymer chain entanglement and storage of energy in the contacts.

As intracellular rheological properties might not only stem from viscous solutes or rigid filaments alone, but also from the interaction of the cytosol with network elements of the cytoskeleton, we asked if Rheo-FLUCS is capable of accurately measuring phase angles of poroelastic materials. For this, we introduced increasing fractions of pre-polymerised linear PAAm chains into the porous network of a cross-linked PAAm gel. While usually higher overall acrylamide concentrations have the tendency to form stiffer and more solid gels, the inclusion of linear chains into the network increases the viscous character of the gel due to strong frictional interaction between the solvent and the gel matrix.^33^ In these experiments, we find that the fluidity significantly increases due to these emergent poroelastic effects that can equally be measured by Rheo-FLUCS.

Figure 3 presents a master curve, combining material information over the entire viscoelastic range spanned by the model PAAm systems. We observed the same quantitative phase angles in these physically different systems across the entire viscoelastic range (*R*^2^ ∼ 0.997, when fitting the means of repeated experiments for each condition). Moreover, we found the uncertainty via Rheo-FLUCS to be lower than that in the magnetic reference measurements, which in parts is likely due to the reduced drift of tracer particles in Rheo-FLUCS compared to the motion of force-subjected probes.

The obtained validation demonstrates the phase-equivalence hypothesis between force-driven and flow-driven microrheology, at the same time confirming that both measurements fall within the linear viscoelastic regime. Through leveraging the contactless and non-invasive nature of localised flow manipulations, we next applied Rheo-FLUCS in living systems.

### Non-invasive intracellular rheology and sensitivity to drug treatment

To test the applicability of Rheo-FLUCS in living cells and its potential as a probe-free technique, we started by using common NIH-3T3 adherent fibroblasts, in which we labelled lysosomes with LysoView 488 as fluorescent tracers (Figure 4a). We find that with labelling these endogenous tracers (tens to hundreds per cell), we bring the sensitivity down to 30 nm oscillation amplitude per lysosome, resulting in phase noise in the range of 10°. Moreover, the simultaneously applied oscillations enable multiplexed readout, which in spite of the large heterogeneity, ensures high statistics.

**Fig. 4.**
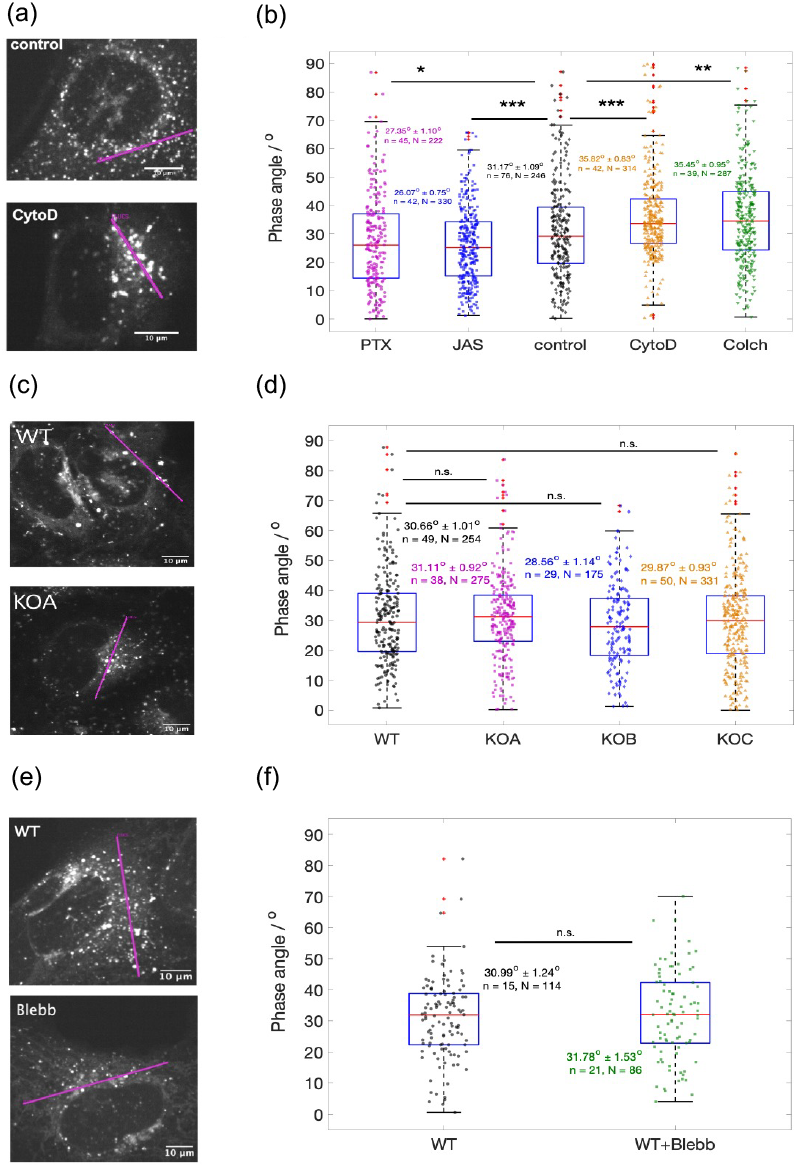
Active probe-free microrheology in compound-treated mouse fibroblasts (NIH-3T3, a-b), precision knockout of myosin isoforms in mammalian osteoblasts (U2OS, c-d) and Blebbistatin-treated mammalian osteoblasts (U2OS, e-f). (a) Fluorescent micrographs of wild-type and drug-treated NIH-3T3 cells (scale bars = 10 *μ*m). The Rheo-FLUCS flows were applied along the magenta lines. (b) Box plot of phase angles measured via Rheo-FLUCS at 1 Hz. Lysosomes labelled with LysoView 488 served as tracers. In each case, we reported the weighted mean with its standard error, number of cells (‘n’), and number of oscillated lysosomes (‘N’) from which information was extracted. In total, we analysed 76 control cells, 45 Taxol-treated (1 *μ*M PTX) cells, 42 Jasplakinolide-treated (1 *μ*M JAS) cells, 42 Cytochalasin-D-treated (1 *μ*M CytoD) cells, and 39 Colchicine-treated (2.5 *μ*M Colch) cells, where treatment was applied 30 minutes prior to the acquisitions. All results were statistically significant when compared with the wildtype (* for p < 0.05, ** for p < 0.01, *** for p < 0.001 as extracted from a two-sample t-test). Red crosses mark outliers. (c) Fluorescent micrographs of wild-type (WT) and Blebbistatin-treated (50 *μ*M Blebb) U2OS cells (scale bars = 10 *μ*m). As in (a), the Rheo-FLUCS flows were applied along the magenta lines. (d) Box plot of phase angles measured via Rheo-FLUCS at 1 Hz. As with NIH-3T3, lysosomes labelled with LysoView 488 served as tracers. In total, we analysed 15 wild-type and 21 drug-treated osteoblasts. No statistical significance (‘n.s.’) was found after applying a two-sample t-test. Red crosses mark outliers. (e) Fluorescent micrographs of wild-type and genetically modified U2OS cells (scale bars = 10 *μ*m), where the NM IIA isoform was genetically deactivated. The Rheo-FLUCS flows were applied along the magenta lines. (f) Box plot of phase angles measured via Rheo-FLUCS at 1 Hz. As with NIH-3T3, lysosomes labelled with LysoView 488 served as tracers. In total, we analysed 49 wild-type (WT) osteoblasts, 38 cells with NM IIA knockout (KOA), 29 cells with NM IIB knockout (KOB), and 50 cells with NM IIC knockout (KOC). No statistical significance (‘n.s.’) was found on applying two-sample t-tests. Red crosses mark outliers.

Next, we asked if the method is suitable to detect changes in cell rheology, for which we used wild-type fibroblasts as well as such treated with compounds known to mechanically stiffen or soften cells. Specifically, we applied drugs employed in chemotherapy that cause alterations in the actomyosin cytoskeleton. This served as a positive control for probing differential mechanics.

First, we applied Paclitaxel (1 *μ*M for 30 min) that is known to stabilise microtubules to the extent of preventing mitosis.^34,35^ In these compound-treated NIH-3T3 cells, we measured lower phase angles relative to the wild type, consistent with reduced cytosol fluidity (Figure 4b). We obtained similar results on treating the mouse fibroblasts with Jasplakinolide (1 *μ*M for 30 min), which was previously reported to promote actin polymerisation *in-vitro*, leading to stiffening of the cytoplasm through strengthening the cytoskeleton.^36,37^ These findings are in agreement with previous reports from microrheology based on atomic force microscopy, where a predominantly elastic response and relative increase in stiffness were observed by Laudadio *et al*.^40^ upon treating rat airway smooth muscle cells with Jasplakinolide.

In contrast, we found upon applying Colchicine (2.5 *μ*M) that suppressing polymerisation of microtubules leads to fluidisation of the cytoplasm, as indicated by the higher phase angles in Figure 4b.^38,39^ Similar effect to the cytosol mechanics we measured upon applying Cytochalasin D (1 *μ*M). The latter can be attributed to capped actin filaments that prevent further polymerisation and, as with Colchicine, effectively weakens the cytoskeleton.^41–43^

We conclude from these measurements that Rheo-FLUCS is capable of detecting changes in adherent cells as small as 4° in the phase angle, which is approximately the difference observed when adding 1%vol linear PAAm to water. Despite the sample-based variability, the multiplexing yields powerful statistics.

### Osteoblast cytoplasm mechanics is insensitive to myosin activity

Motivated by recent findings obtained with a cell microstretcher,^44^ showing that cortex mechanics is finely controlled through a complementary interplay of three nonmuscle myosin II (NM II) isoforms (A, B and C), we asked if the mechanical properties of the cytoplasm are similarly dependent on motor activity as the cortex. Previously, it was suggested that NM IIA is essential for building up cellular tension during initial stages of force generation, whereas NM IIB is required to elastically stabilise NM IIA-generated tension. On the other hand, a new role of NM IIC in establishing tensional homeostasis has been revealed.^44^

To investigate further the individual contributions of each NM II isoform to cytosol mechanics, we performed NM IIA, NM IIB and NM IIC CRISPR/Cas-based genetic knockouts in U2OS mammalian osteoblasts.^45,46^ In each case we did not detect any observable change compared to the wildtype (Figures 4c-d), thereby concluding that, although NM II isoforms play a major role in establishing a fine balance of mechanical forces in the cortex, individually they do not significantly contribute to cytosol stiffness.

As none of the motor knockouts led to substantial changes of cytoplasm mechanics individually, we asked if NM isoforms could still have redundant functions with respect to cytoplasm mechanics. Therefore, we applied a compound treatment (50 *μ*M Blebbistatin, Figures 4e-f) in the same mammalian U2OS cell line. Applying Blebbistatin drug treatment to suppress all myosin isoforms simultaneously resulted in no net effect on cytosol mechanics, emphasising that the isoforms neither have individual roles with regards to cytoplasm mechanics, nor do they have redundant roles. This observation was further corroborated by applying 50 *μ*M Blebbistatin treatment in NIH-3T3 cell line (Supplementary Figure 1).

Comparing with the work of Weißenbruch *et al*.,^44^ we find a disparate response in cytosol and cortical mechanics, which seems unexpected since the actomyosin cytoskeleton has been identified as the canonical regulator of cell shape. Rapid cell changes induced by anisotropic cortical contractions have a profound impact on the cytosolic fluid dynamics during cell migration and cytokinesis.^47–49^ However, although inhibition of actomyosin contractility was shown to reverse cytosolic flow direction in keratocytes,^47^ our results suggest that cortical rigidity does not intrinsically influence the stiffness of the cytosol in adherent mesenchymal cells.

The Rheo-FLUCS results emphasise the importance of deciphering the mechanisms responsible for cortex-cytosol mechanical interplay that appear to reach beyond a simple description of actomyosin cytoskeleton – membrane coupling.

### Applying FLUCS microrheology to ascidian oocytes

Ooplasm mechanics play a pivotal role in development within biological systems and hold significant promise for applications in reproductive medicine.^50–52^ For example, in mouse oocytes, asymmetric streaming crucially positions the spindle complex, underscoring the intricate choreography involved in embryonic development.^53,54^ Moreover, friction forces are instrumental in dictating cytoplasmic reorganisation, positioning material and inducing shape changes in ascidian oocytes upon fertilisation.^6^ Given the contribution of the mechanical properties of oocytes to embryonic development, we asked whether we could measure ooplasmic properties of ascidians, a simple yet powerful model organism of the large phylum *Chordata*, with an egg of around 150-*μ*m diameter, using Rheo-FLUCS.

To achieve non-invasive measurements, we labelled yolk granules in *Phallusia mammillata* oocytes with SYTO 59 and employed Rheo-FLUCS to oscillate them instead of microbeads (Figure 5a and Supplementary Figure 3). Due to the high density of yolk granules, these measurements could be highly parallelised. By targeting thousands of yolk granules per confocal section in each oocyte, we achieved comprehensive mechanical profiling across multiple positions simultaneously.

**Fig. 5.**
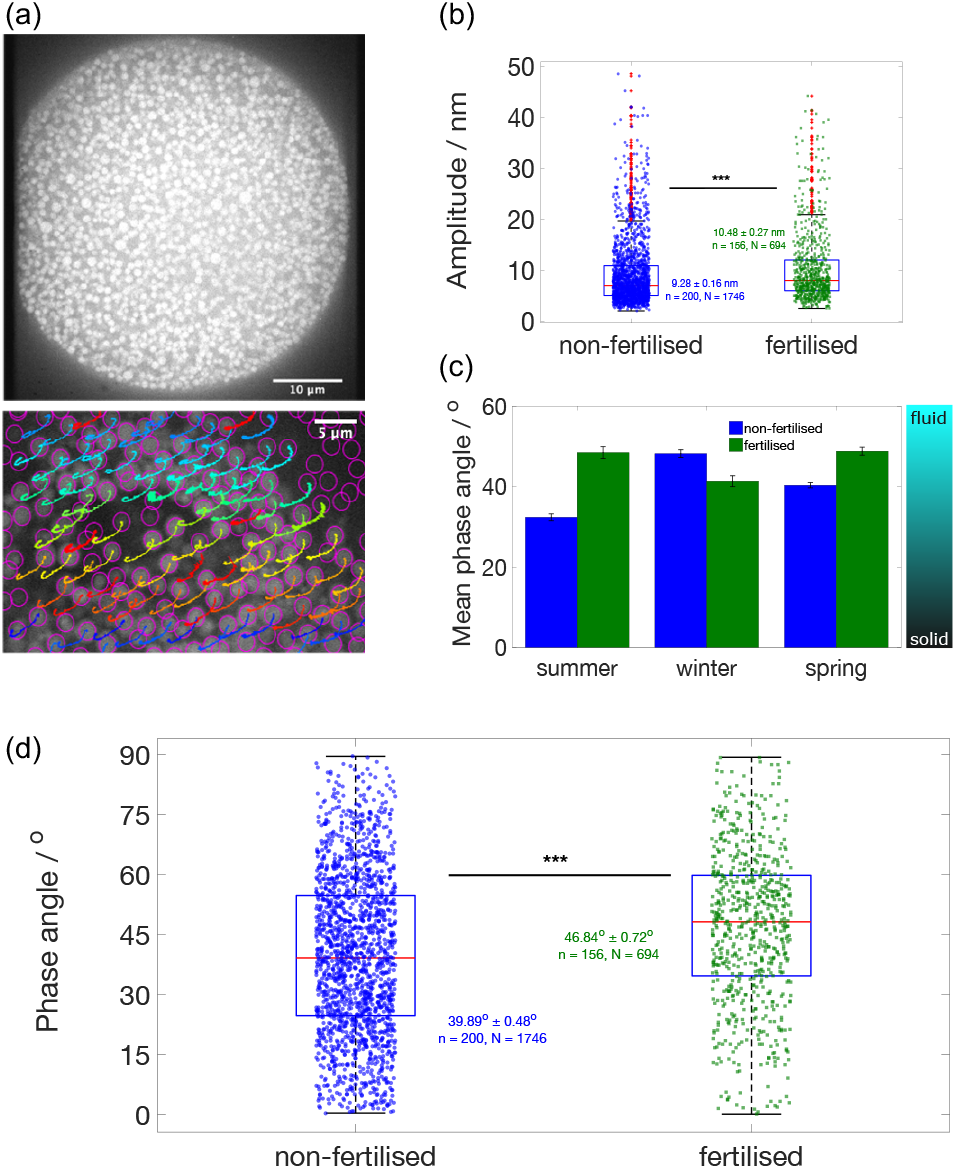
Mechanosensing in non-fertilised and fertilised ascidian oocytes (*Phallusia mammillata*). (a) SYTO-59 labelling of yolk granules within a dechorionated ascidian oocyte (scale bar = 10 *μ*m). The granules were simultaneously oscillated via Rheo-FLUCS at 1 Hz frequency and then tracked via the TrackMate plugin on Fiji (scale bar = 5 *μ*m). (b) Amplitude box plot generated from the oscillatory motion of yolk granules in non-fertilised and fertilised (first-polar-body stage) ascidian eggs. (c) Mean phase-angle data collected across three different seasons, displaying variations in the mechanical response of both non-fertilised and fertilised oocytes. Each error bar represents standard error of the mean, where the data were collected from a total of 20 non-fertilised and 15 fertilised eggs in the summer, 65 non-fertilised and 45 fertilised eggs in the winter, and 115 non-fertilised and 96 fertilised eggs in the spring. (d) Phase-angle box plots, showing the overall increase in phase angle upon fertilisation across all three seasons. In both (b) and (d) we report the distribution’s mean and standard error of the mean, number of analysed oocytes (‘n’), and number of oscillated yolk granules (‘N’). The differences in both cases were found to be statistically significant (p < 0.001) on performing a two-sample t-test.

In this work we encountered several factors that influence ooplasm mechanics, contributing to better understanding of how these dynamics are regulated within ascidian oocytes. First, we analysed the motion of the yolk granules and extracted the amplitudes (Figure 5b) and phase angles (Figures 5c-d) of oscillations induced by Rheo-FLUCS. We compared those prior to and moments after fertilisation. The reference measurement point in all fertilisation experiments was the ejection of the first polar body during meiosis. Overall, fertilised oocytes exhibited increased fluidity compared to their non-fertilised counterparts. This finding aligns with existing reports indicating that active processes initiated by fertilisation enhance the fluid dynamics of the ooplasm, thereby facilitating subsequent developmental stages.^55–58^ The increase in fluidity likely plays a crucial role in enabling cellular rearrangements necessary for successful embryonic progression.

Further, our research identified significant seasonal variations in ooplasm rheology. These findings complement existing literature that highlights the seasonality associated with reproductive potential in various species.^59–61^ This highlights the promising use of Rheo-FLUCS in the field of reproductive medicine for predicting or evaluating reproductive capacity based on rheological properties. Ooplasm rheology could thus serve as a window into the intricate processes governing early development, with the outlook of advancing fertility treatments and improving conservation efforts aimed at preserving biodiversity.

## Discussion

In this work, we demonstrated that intracellular active microrheology is possible without the need for an internalised force probe. By establishing the equivalence of flow- and force-based phase angles across a wide range of materials, with fundamentally different microscopic properties underlying their emergent rheological behaviors (liquids, viscoelastic fluids, poroelastic gels, and solid gels), we enable canonical rheological measurements even in living systems, from single cells to early embryos.

The technology opens new avenues for exploration in materials science and mechanobiology, where direct access to mechanics was previously limited due to compartmentalisation and probe incompatibility. The contactless flows induced by Rheo-FLUCS both extend the measurable viscoelastic range of complex fluids and enable readout of the mechanical state of cellular cytoplasm and embryonic ooplasm. While Brillouin spectroscopy informs about mechanical properties at gigahertz frequencies,^21–24^ where thermal diffusion dominates, our technique extends non-invasive measurements well below 10-100 Hz, where active processes are known to have an influence on the net mechanical properties. Successive phases of diffusion and active transport have been shown through temporal analysis of single-particle transport on trajectories of labeled vesicles^62,63^ or viruses^64^ in single cells, where path dissection was performed manually.^65^ Hence, measuring cell mechanics on timescales longer than 10 ms necessitates the use of active methods discriminating between passive and active motion.^66,67^

Finally, this opens possibilities to use Rheo-FLUCS in the detection of physiological changes in cellular phenotypes, *e*.*g*. as was previously shown via the cell’s metabolic state,^14^ and gain additional power and universality by expanding the readout into the time domain. Within the life sciences, performing Rheo-FLUCS could enable uses of probe-free rheology as a diagnostic tool for disease and reproductive medicine.^63^

## Methods

### FLUCS interactive microscopy setup

All experiments were conducted using the Focussed Light-Induced Cytoplasmic Streaming (FLUCS) microscope described in our previous work,^18^ where simultaneous external triggers were sent synchronously to the laser scanner (two-axis acousto-optic deflector, model AA.DTSXY-A6, Pegasus Optik) and the imaging device (Zyla 5.5 sCMOS camera). The infrared laser beam (1,455 nm wavelength, Raman laser, CRFL-20-1455-OM1, 20 W, Keopsys, CW mode) was focussed through a custom-coated 60× objective lens (UPLSAPO, NA = 1.2, W-IR coating, Olympus) and was then used to generate dynamic heat patterns. The laser scanning in all microrheological experiments was performed at 2 kHz scan rate, where the laser was repeatedly scanned 1,000 times along each of two opposite directions. Since the direction of the induced flows is determined by the direction of laser scanning, the flows were reversed every 0.5 seconds. This effectively induced steady-shear oscillations with a period of close to 1 second, which contains an approximate error of 600 *μ*s due to signal processing.

### Software control of the FLUCS setup

Custom-written LabView software was used to generate the waveforms required for scanning the laser via the acousto-optic deflector. A data acquisition card (*cf*. Mittasch *et al*.^14^) served as an analog-to-digital converter and enabled rapid signal transfer between the computer and the laser scanner. The frequency and transition of the waveforms were verified using an oscilloscope (Datatec, HMO72). The LabView software allowed real-time modifications to the frequency of laser scanning, linear extent of the flows, number of scan repetitions, duration of the acquisition and other experimental parameters.

### Software for analysing active, phase-sensitive microrheology data

Imaging was performed by Zyla 5.5 sCMOS camera and the motion of probe-particles tracked via Fiji’s open-source TrackMate plugin.^68^ Subsequently, the tracked particle coordinates and temporal data were forwarded to an in-house Python routine that served as a monochromatic filter that selects viscoelastic data at the frequency of oscillation. Most notably, the routine allowed decomposition of the filtered signal into an amplitude and a phase, where the latter served as a precise measure of material properties. The generation of box plots and statistical analysis were performed on Matlab.

### Linear and cross-linked polyacrylamide preparation

We followed the protocol for preparing polyacrylamide gels outlined by Charrier *et al*.^33^ Linear polyacrylamide solutions (50 mL volume) were prepared in the absence of a cross-linker by mixing 40% acrylamide (Bio-Rad, #1610140) with N, N, N’, N’-tetramethylethylenediamine (25 *μ*L TEMED, Sigma-Aldrich, T9281) and 10% ammonium persulfate (120 *μ*L APS, Sigma-Aldrich, A3678). The mixture was then degassed using a vacuum desiccator for 30 minutes and incubated overnight at 37 °C. The resulting highly viscous solution was combined in different proportions with a cross-linked polyacrylamide gel prepared using 40% acrylamide, TEMED, APS and 2% bis-acrylamide (Bio-Rad, #1610142). The final acrylamide (monomer-to-bis) ratio was 100:1.

The probe-particles we used in the flow-based microrheology experiments on polyacrylamide gels were fluorescent (Dragon Green) 1.037-*μ*m carboxylated polystyrene beads (PS-COOH uniform dyed microspheres, Bangs Laboratories, Inc., FC04F). These were diluted 10^3^× in the test sample and then pipetted into a 15-*μ*m tall chamber composed of a plain microscope slide (76×26×1 mm, Paul Marienfeld GmbH & Co. KG, Germany) and a thin round cover slip (22-mm diameter, 170-*μ*m thickness, borosilicate glass). The chambers were sealed with two-part impression material (Vinylsiloxanether, Identium, 13711) to minimise evaporation over the course of the measurements.

### Magnetic microrheology validation measurements

For performing magnetic-force driven microrheology experiments, we used electromagnetic needle as reported in Stoev *et al*.^18^ The needle was of 4.5 mm diameter, 100 mm length and the core of the electromagnet was Hy-Mu 80 (80% nickel-iron-molybdenum alloy, rod dimensions of 0.380×0.135 in, National Electronic Alloys Inc., Oakland, NJ), enclosed within a brass frame. To ensure the induced magnetic forces were sufficiently strong to oscillate particles within polyacrylamide gels, we modified the tip angle to 30° and the number of coil turns was increased to 650. We performed the experiments in an open-chamber configuration, where the sample was loaded onto a customised MatTek dish with a wall opening for the needle. The latter was connected to a laboratory DC power supply (EA Elektro Automatik, EA-PS 3016-10B) via LED driver that allowed periodic on-off pulsing of the magnetic field. The probe-particles we employed for these measurements were fluorescent (Dragon Green) 0.9-*μ*m magnetic polymer-coated beads (PS / 6% Divinylbenzene / V-COOH Mag 480-520, Bangs Laboratories, Inc., MCDG001) and 8.26-*μ*m magnetic COMPEL beads (COOH-modified, 480-520, Bangs Laboratories, Inc., UMDG003).

### Preparation of NIH-3T3 cells for FLUCS microrheology

Mouse embryonic fibroblasts (NIH-3T3) were cultured in DMEM high glucose (Gibco, 31966-021) supplemented with 10% fetal bovine serum (Gibco, 10270-105) and 1 µg/mL penicillin and streptomycin (Gibco, 15140-122) in a humidified incubator with 5% CO_2_ at 37°C.

For the microrheology experiments, the cells were seeded onto a clean glass coverslip and allowed to reach approximately 50% confluency. On the day of the FLUCS experiment, 30 min in advance the cells were treated with compounds, *viz*. 2.5 µM Colchicine (Sigma-Aldrich, C3915), 1 µM Paclitaxel (Sigma-Aldrich, T7191), 1 µM Jasplakinolide (Sigma-Aldrich, J4580) and 1 µM Cytochalasin D (Sigma-Aldrich, C8273), and co-stained with LysoView™ 488 (1:800 dilution, Biotium, 70067). After the treatment, the cells were carefully loaded onto an experimental chamber made of a thick sapphire glass slide with Peltier elements on each side to control the temperature, as described by Mittasch *et al*.^14^ Polystyrene beads (15-µm diameter) dispersed 1,500× in culturing medium were added as a spacer between a coverslip and a microscope slide to prevent excessive compression of the cells during the experiments.

The FLUCS microrheology experiments were then conducted by generating linear oscillatory scans (1 Hz frequency) in the cytoplasm of the cells and away from the nucleus. The stained lysosomes were used as tracers and their motion subsequently analysed using the same particle tracker and Python code as with the synthetic probe-particles in the polyacrylamide gel experiments. Only phase lags with uncertainty less than 30° were included in the final plot.

### Preparation of U2OS osteoblasts for FLUCS microrheology

Human osteosarcoma cell line (U2OS), wild-type and CRISPR knockouts for the nonmuscle myosin II (NM II) isoforms A, B and C genes, were used to determine the effect of each gene on tensional homeostasis. As with NIH-3T3, U2OS cell lines were cultured in DMEM (Gibco, 31966-021) supplemented with 10% FBS (Gibco, 10270-105), 1ug/mL Penicillin streptomycin (Gibco, 15140-122). The cells were maintained in a humidified 5% CO_2_ incubator at 37 °C. For the Rheo-FLUCS experiments, the cells were seeded onto a glass coverslip to reach 50-60% confluency. On the day of the experiment, the cells were treated with LysoView™ 488 (1:800 dilution, Biotium, 70067) for 30 minutes. The coverslips were then carefully loaded onto an experimental chamber made of a thick sapphire glass slide with Peltier elements on each side to control the temperature, as described by Mittasch *et al*.^16^ Polystyrene beads (15-µm diameter) dispersed 1,500× in culturing media were added as a spacer between a coverslip and a microscope slide to prevent excessive compression of the cells and preserve the viability of the cells over the course of the experiments.

### Preparation of ascidian oocytes for FLUCS microrheology

*Phallusia mammillata* were obtained from Roscoff Marine Station (EMBRC, France) and kept in artificial seawater (ASW; 36 g/L Tropic Marin® BIO-ACTIF Sea Salt, Tropic Marine) at 16°C for 3-4 weeks under constant light. Eggs and sperm were harvested and kept at 4°C for 3-4 days. Eggs were dechorionated with 1% trypsin (Sigma-Aldrich, T8003) in ASW for 40 minutes under gentle agitation and kept in ASW supplemented with 0.05 g/L streptomycin sulfate salt (Sigma-Aldrich, S6501). Dechorionated and non-fertilised eggs were incubated with 1 µM SYTO 59 red fluorescent nucleic acid stain (Invitrogen, S11341) in ASW for 5 min and washed with ASW prior to fertilisation. Ascidian sperm was activated with pH 9 ASW and used to fertilise dechorionated eggs. All imaging and FLUCS experiments were conducted at 15°C.

We performed Rheo-FLUCS experiments and analysis as with the NIH-3T3 cells, where in the ascidian eggs we generated linear oscillatory scans deep into the ooplasm and away from the cell membrane. These were again conducted at 1 Hz oscillation frequency, at which we compared the response of non-fertilised and fertilised eggs.

## Data and code availability

Source data and code are available from the corresponding author upon reasonable request.

## Acknowledgements

IDS and MK kindly acknowledge funding from the Life grant by Volkswagen Foundation (Life! Grant No. 92772), Deutsche Forschungsgemeinschaft (DFG, German Research Foundation, Germany’s Excellence Strategy, 2082/1, Grant No. 515462906) and the Hector Foundation. IDS was additionally funded by the Karlsruhe Institute of Technology Excellence Strategy via the Young Investigator Group Preparation Program. VRK, SW and MK thank the ERC Starting Grant GHOSTs (Grant No. 853619). The authors gratefully acknowledge help from Dr. Benjamin Seelbinder, Mr. Claudius George and Mr. Falk Elsner in designing and constructing the magnetic needle for force-driven rheology experiments. We thank Dr. Iain Patten for valuable discussions on the structure of the manuscript and Ivan Saraev for graphics support.

## Author information

*Max Planck Institute of Molecular Cell Biology and Genetics, Dresden, Germany* Iliya D. Stoev, Venkat R. Krishnaswamy, Susan Wagner, Elena Erben & Moritz Kreysing

*Institute of Biological and Chemical Systems - Biological Information Processing (IBCS-BIP), Karlsruhe Institute of Technology, Karlsruhe, Germany*

Iliya D. Stoev, Venkat R. Krishnaswamy, Susan Wagner, Elena Erben, Martin Bastmeyer & Moritz Kreysing

*Institute of Science and Technology Austria, Klosterneuburg, Austria* Madison Bolger-Munro & Carl-Philipp Heisenberg

*Department of Neuroscience, Universitá Catolica del Sacro Cuore, Rome, Italy* Antonio Minopoli

*Department of Cell and Developmental Biology, University College London, London, UK* Kai Weißenbruch

*Zoological Institute, Cell and Neurobiology, Karlsruhe Institute of Technology, Karlsruhe, Germany*

Kai Weißenbruch & Martin Bastmeyer

*Human Technopole, Milan, Italy*

Nicola Maghelli

### Contributions

IDS and MK conceived the project. IDS, MBM and VRK conducted the experiments. IDS analysed the data. MK provided the analysis code, which EE and AM further developed. NM enabled hardware implementation (laser-camera synchronisation). IDS wrote the first draft, which IDS and MK finalised. VRK and SW performed cell culture, genotyping and preparation of cells for experiments. KW and MB participated in valuable discussions about Rheo-FLUCS data on U2OS cells. MBM and CP provided valuable input for conception and analysis of ascidian microrheology data. All authors participated in critical discussions of the final draft.

## Ethics declarations

### Competing interests

IDS and MK applied for an international European patent for force measurement technology related to this publication (application number PCT/EP2021/071392). MK is a co-inventor of technology for laser-induced flow and optofluidic technology (EP3714308A1, US patent number 17/506,750 and PCT/EP2021/071437) and holds a consultancy contract with Rapp OptoElectronic GmbH.

