## Supplementary Information for "Active and Probe-Free Intracellular Rheology via Phase-Sensitive Thermoviscous Flows"

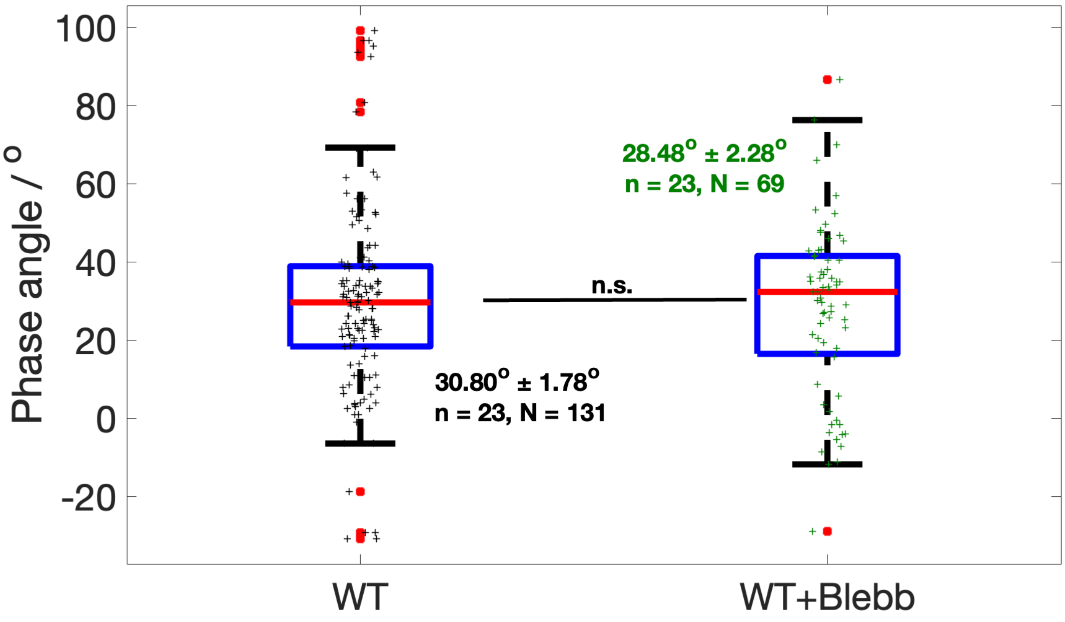


**Supplementary Fig.1 Mechanosensing in NIH-3T3 mouse fibroblasts on Blebbistatin treatment.**

Phase-angle box plots, showing that the cytosol of NIH-3T3 mouse fibroblasts remains unchanged after Blebbistatin drug treatment. Here we include weighted mean of the distribution, standard error on the weighted mean, number of analysed cells ('n'), and number of oscillated lysosomes ('N').

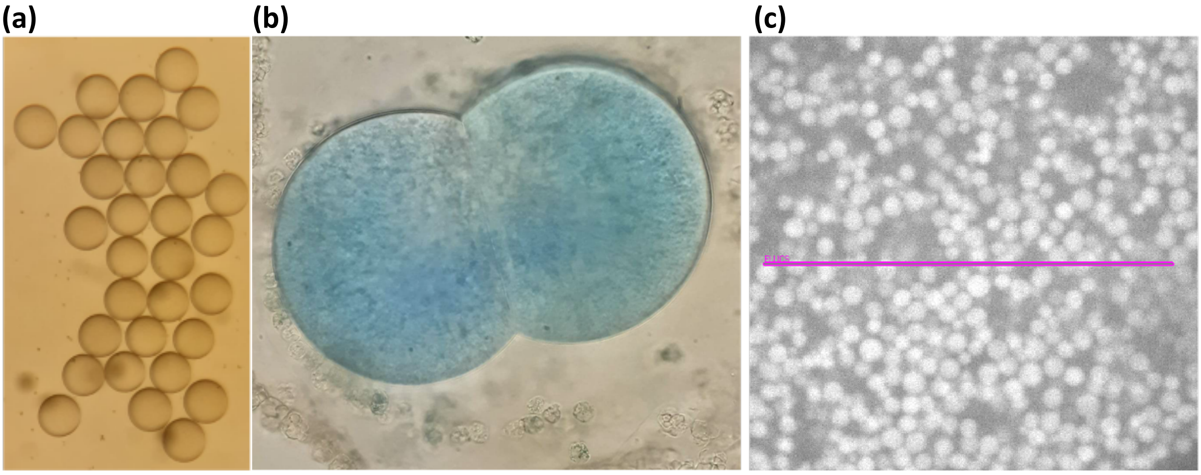


**Supplementary Fig. 2 Dechorionation, staining and scanning of ascidian oocytes.**

(a) Adding a small amount of trypsin was found to successfully remove the chorion of the ascidian oocytes. The exact concentration and volume of added trypsin strongly depended on the number of eggs that were dechorionated. Smaller volumes of trypsin generally require longer incubation times for complete dechorionation.

(b) SYTO-59 staining of the yolk granules and Rheo-FLUCS laser scanning did not interfere with the natural course of division of ascidian oocytes which were fertilised following the addition of the dye and divided into two identical blastomeres.

(c) Example of a Rheo-FLUCS scan line that simultaneously oscillated tens of yolk granules, thus extracting mechanical information from multiple regions within the ooplasm. The displayed yolk granules vary in size between 10 and 20 microns.

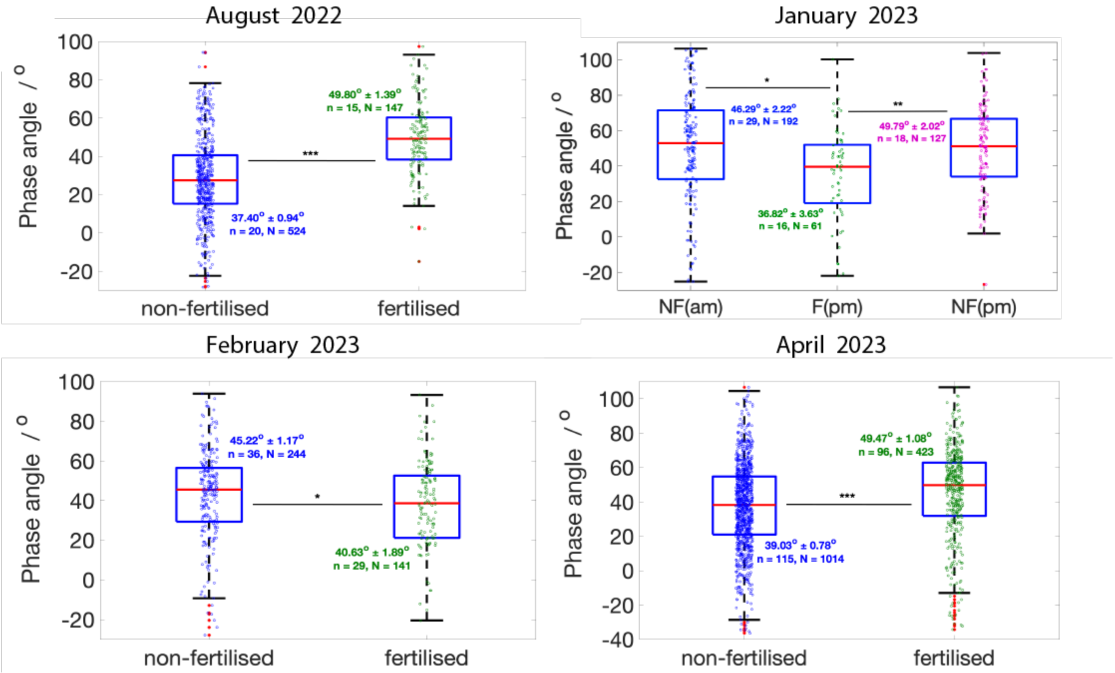


**Supplementary Fig.3 Mechanosensing in ascidian oocytes over four different months.**

Phase-angle box plots, showing the increase in phase angle upon fertilisation in spring/summer (August 2022 and April 2023), and drop in the phase angle after fertilisation in the winter (January and February 2023). We show information about the weighted mean of the distribution, standard error on that weighted mean, number of analysed oocytes (‘n’), and number of oscillated yolk granules (‘N’).

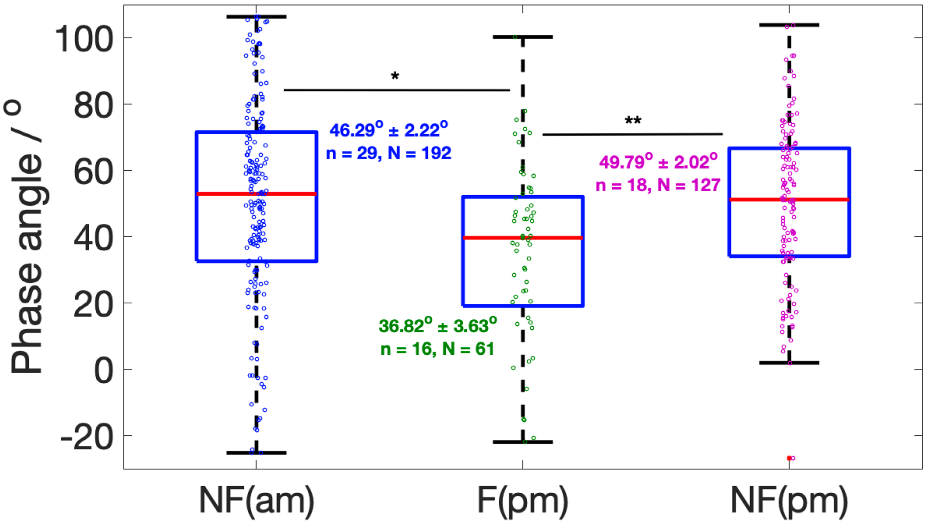


**Supplementary Fig.4 Phase-angle variations in the ooplasm of ascidian oocytes during the month of January.**

Phase-angle box plot, showing the decrease in phase angle upon fertilisation in the month of January. This was in contrast to the data obtained from the spring and summer months. However, there was no significant variation in measuring non-fertilised (NF) oocytes within a few hours, indicating that the mechanical information remains relatively constant throughout the day. We show information about the weighted mean of the distribution, standard error on that weighted mean, number of analysed oocytes ('n'), and number of oscillated yolk granules ('N').

(a) Magnetic Needle Pulling

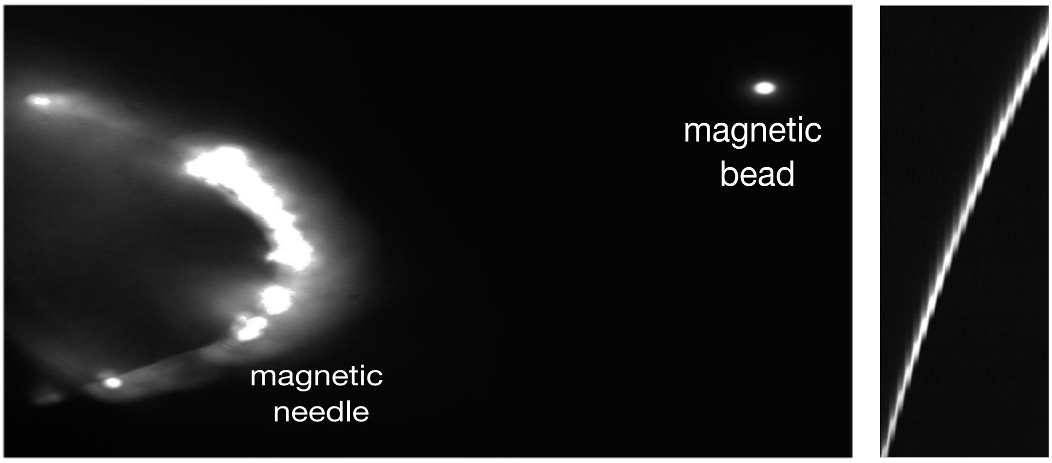


(b) Fourier Transform Analysis

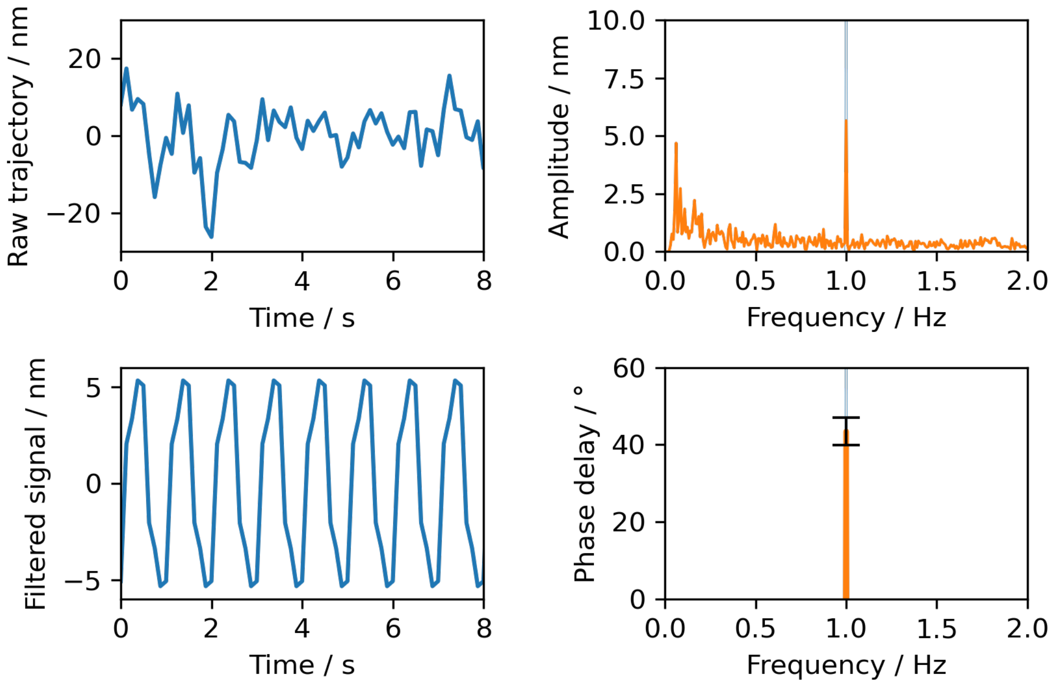


**Supplementary Fig.5 Reference microrheology measurements requiring the application of external magnetic forces and the internalisation of a magnetic probe.**

(a) A pulsed current was applied to an electromagnetic needle which periodically pulled on a magnetic bead. The fluorescent image shows 0.9-$\mu$m magnetic bead approaching the needle, where we visualised the oscillations via a kymograph.
(b) As with the flow-based microrheology, Fourier-transform analysis was used to convert the raw oscillatory trajectory of the magnetic probe into the mechanical response of the material. Filtering allowed clean extraction of the phase delay between the driving stimulus and the sample response, given a known oscillatory frequency, here 1 Hz. The displayed data are for the same

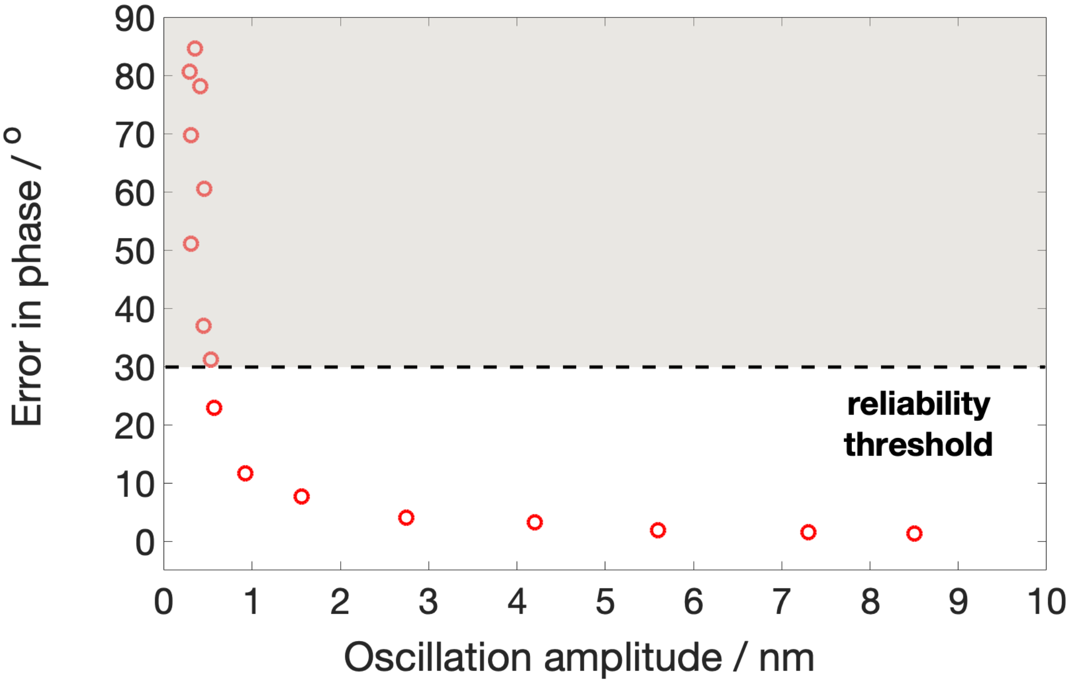


**Supplementary Fig.6 Rheo-FLUCS limit of detection in a highly viscous tar pitch.**

Rheo-FLUCS can be applied in media of arbitrary viscosity and this is demonstrated here with a very highly viscous tar pitch, where the amplitude of the oscillations directly depends on the laser power. As expected, we observed an increase in the uncertainty of the phase lag on lowering the laser power, where remarkably even at sub-nanometer oscillation amplitudes, we could determine phases with a precision of 12°. We did not include in the manuscript any data with error in the phase lag of over 30°.

**Supplementary videos 1-3:** Bright-field and fluorescent tracking of the progression towards division of a single *Phallusia mammillata* fertilised egg in the spring/summer months.

**Supplementary videos 4-6:** Bright-field and fluorescent tracking of the progression towards division of a single *Phallusia mammillata* fertilised egg in the winter months.

**Supplementary video 7:** Rheo-FLUCS oscillations induced in a Newtonian sample using a 1-$\mu$m bead.

**Supplementary video 8:** Magnetic-tweezer microrheology, showing a creep experiment, where the magnetic field induced in an electromagnetic needle was pulsed periodically, leading to pull-release cycles. The magnetic bead here was of size 1-$\mu$m.

**Supplementary video 9:** Rheo-FLUCS oscillations within a highly viscous tar pitch, similar to the one used in the famous pitch-drop experiment. This demonstrates the utility of the method in highly viscous media, where passive microrheology techniques relying on Brownian motion fail to perform.

**Supplementary video 10:** Rheo-FLUCS application in bright field, where a 1-$\mu$m bead is oscillated within an artificial cell composed of intralipid and agarose.

**Supplementary video 11:** Rheo-FLUCS application in bright field, where test cells and follicle cells in chorionated ascidian eggs are oscillated. This particular egg was not selected for further analysis due to its inability to fertilise.
